# Ventral premotor cortex influences spinal cord activation during force generation

**DOI:** 10.1101/2023.02.22.529375

**Authors:** Hanna Braaß, Jan Feldheim, Ying Chu, Alexandra Tinnermann, Jürgen Finsterbusch, Christian Büchel, Robert Schulz, Christian Gerloff

**Affiliations:** Department of Neurology, University Medical Center Hamburg-Eppendorf, 20246 Hamburg, Germany; Institute of Systems Neuroscience, University Medical Center Hamburg-Eppendorf, 20246 Hamburg, Germany

## Abstract

Force generation is a crucial element of dexterity and a highly relevant skill of the human motor system. How cerebral and spinal components interact and how spinal activation is influenced by cerebral primary motor and premotor areas is poorly understood. Here we conducted combined cortico-spinal functional MRI during a simple visually guided isometric force generation task in a group of 20 healthy young subjects. Activation was localized in the ipsilateral cervical spinal cord and contralateral primary motor and premotor areas. The main finding is that spinal activation was influenced by ventral premotor cortex activation. Spinal activation was furthermore significantly correlated with primary motor cortex activation while increasing target forces led to an increase in the amount of activation. These data indicate that human premotor areas such as the ventral premotor cortex might be functionally connected to the lower cervical spinal cord contributing to distal upper limb functions, a finding which extends our understanding about human motor function beyond the animal literature.

## Introduction

Force generation and modulation are highly relevant prerequisites of human hand dexterity and important for mastering activities of daily living. Seminal neuroimaging studies ^1–7^, studies in patient cohorts ^8–10^ and systematic reviews ^11,12^ have significantly contributed to our current understanding of cortical and subcortical representations of force generation and control. Key nodes of this network comprise the primary motor cortex (M1) and multiple secondary motor cortices of the frontal ^13–16^ and parietal lobe ^17^, basal ganglia ^18^, and the cerebellum ^19^. Key pathways primarily include the cortico-spinal tract (CST), moreover cortico-cortical tracts, and various cortico-fugal pathways between the cortex, basal ganglia, and the cerebellum.

Technical advantages in neuroimaging sequences ^20–22^, denoising ^23^, and spinal analysis ^24^ have recently paved the way for combined cortico-spinal functional MRI. It has been applied to explore motor network dynamics during rest ^25^ or during complex hand movements ^26^. These studies have extended the prior knowledge derived from a variety of earlier spinal functional MRI studies ^27^. They have added spinal data with a high spatial resolution to previous electrophysiological data of cortico-spinal information throughput, such as those derived from cortico-muscular coherence analyses ^28–31^.

So far, combined cortico-spinal functional imaging during force generation has not been studied systematically. It would provide novel insights into multi-site cortico-spinal interactions during force generation, particularly in the context that not only M1, but also multiple secondary motor areas contribute to the CST and send trajectories to the spinal cord ^32,33^. In detail, studies of non-human primates have provided converging evidence that regions such as the dorsal and ventral premotor cortex (PMV) on the lateral surface of the hemisphere and the supplementary motor area (SMA) on the medial wall show cortico-spinal structural connectivity and have the potential to act in parallel to generate output to the spinal cord ^34–43^. However, even in primates, there are still many open questions regarding potential relay nodes, and the precise craniocaudal extent of mono-or, more likely indirect poly-synaptic connectivity ^44^ along spinal inter-neural routes including the proprio-spinal system ^45–47^. For instance, tracing data in macaques have shown that PMV contributes about 5% of the total cortico-spinal projection with the majority terminating already in the upper segments of the cervical spinal cord, with only few terminating more caudally ^38,39,48^. Data obtained in rhesus monkeys did not show PMV projections below cervical level 2, but prominent terminal medial spinal densities for dorsal premotor cortex between C5 and T1 ^49^. Data in humans are strikingly limited. Transcranial magnetic stimulation (TMS) and multi-regional analysis of motor-evoked potentials have suggested the existence of fast and direct functional cortico-spinal connectivity between dorsal premotor areas and the hand ^50,51^. Using TMS, peripheral nerve stimulation and monosynaptic reflex analysis, another study argued for the existence of descending influences onto proprio-spinal inter-neurons not only from M1 but also from PMV ^52^. Recent tractography data have suggested potential direct connections between premotor areas including PMV and SMA and even lower cervical segments. However, these should be considered with caution due to reduced spatial resolution at the spinal levels ^53^. Finally, studies in stroke patients have related the extent of damage to CSTs originating from premotor areas to deficits and recovery processes ^54–56^.

In summary, there is converging data to hypothesize that spinal cord activation during force generation might be significantly linked to contralateral M1 activation. In addition, and importantly, it can also be hypothesized that the extent of spinal activation might be modulated by premotor regions such as PMV and SMA. The present study was designed to explore these hypotheses in detail. Cortico-spinal functional MRI data were acquired during a simple visually guided isometric force generation task in a group of healthy young subjects. Brain activation was localized in contralateral M1, SMA and PMV and ipsilateral cervical spinal cord. Mixed-effects linear regression analyses were used to compare cortical and spinal activations between different force levels and to relate cortical activations in M1, SMA and PMV to spinal activations.

## Results

### Subjects and motor task

Twenty healthy young subjects (10 females and 10 males, mean age 27 years, range 19-34) were included in the study. The subjects underwent cortico-spinal functional MRI during a grip force experiment. It comprised repetitive, almost isometric hand grips with the right, dominant hand. There were three different target force levels in a block design, which were low, medium, and high with predefined target force levels corresponding to 30%, 50%, 70% of the linear force continuum covered by an MRI compatible grip force response device (Grip Force Bimanual, Current Design, Inc, Philadelphia, PA). Average maximum grip force across the group was 42±10 kg (mean ± standard deviation, range 31.3-71.3 kg). During the experiment, the exerted target forces were 37.1±6.3% for low, 58.4±7.6% for medium and 76.5±5.8% for high, respectively. The actual target force was overachieved by 7.36%, with no significant difference between the three force levels.

### Spinal cord and cortical brain activation during force generation

Force generation across force levels led to a significant BOLD activity on group level primarily in the ipsilateral right spinal cord between the lower parts of the C5 and C7 vertebral level, with a maximum activity localized at C6 vertebral level. This level would correspond to the C7 spinal cord segment (Fig. 1A). Force level specific analyses resulted in an increase in the spatial extent of BOLD responses from rather focal activation at C6 vertebral level during low force generation towards more distributed spinal activations between C5 and C7 and a peak between C6 and C7 (vertebral level) during high force generation. This extent would correspond to lower C7 and upper C8 spinal cord segments (Fig. 1B, see Supp. Tab. S1 for statistics on peak activations and force-dependent increase in spatial activation).

**Figure 1.**
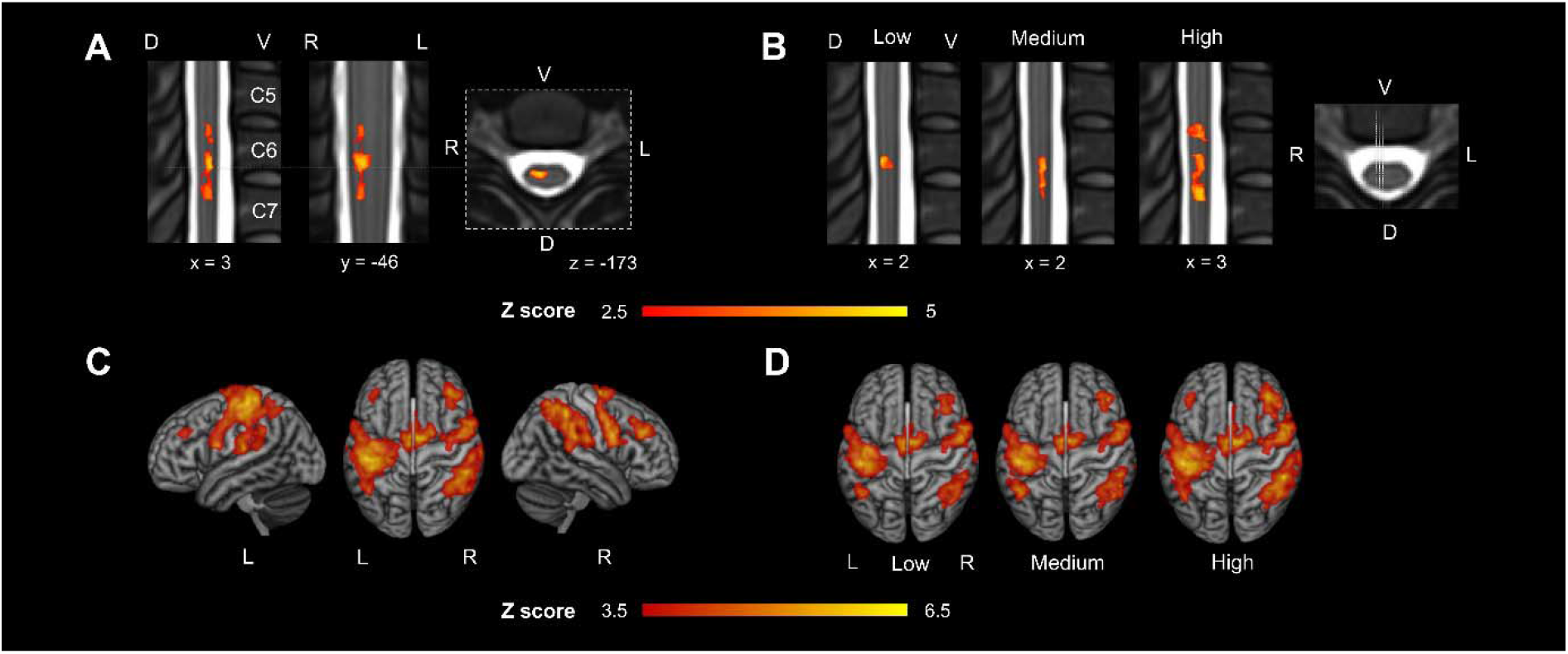
Topography of spinal cord and cortical brain activation during force generation. Estimated group mean spinal BOLD response (Z-maps, thresholded by Z>2.5, cluster significance threshold of *P*<0.05, maximum force level (kg) was included as additional confound parameter) across the three force levels (mean EV, explanatory variable) on two sagittal and one transversal slice in **A**, for low, medium, and high target force level individually on sagittal slices in **B**. Spinal cord activations are overlaid on the PAM50_t2-template. Estimated group mean cerebral BOLD response (Z-maps, thresholded by Z>3.5, cluster significance threshold of *P*<0.05) are plotted for the mean EV across force levels in **C** and individually for each force level in **D**. Cerebral activations are rendered on a T1 template in MNI space. L=Left, R=Right, V=Ventral, D=Dorsal.

Cerebral BOLD activity on group level was detected across force levels primarily in the contralateral primary sensorimotor cortex comprising M1 and the primary sensory cortex (Supp. Tab. S2). We also found activations in bilateral SMA, ipsilateral dorsal premotor cortex, bilateral regions corresponding to PMV and a widespread activation in posterior parietal cortices along the intraparietal sulcus (Fig. 1C, Supp. Tab. S2). Increasing force levels resulted in more pronounced brain activation in contralateral primary sensorimotor cortices and bilateral prefrontal and parietal brain regions (Fig. 1D). The contralateral dorsal premotor cortex did not show significant activations on group level.

In the following, spinal cord and cerebral peaks of activation were localized in each individual subject in the ipsilateral spinal cord and contralateral motor cortices M1, PMV and SMA (see Supp. Tab. S3 for coordinates) using the mean effect contrast across the three force levels (mean explanatory variable, EV). Subsequently, parameter estimates of the BOLD response were extracted on these coordinates for the individual force levels, i.e., three estimates were obtained for all regions in each subject for further statistical analyses.

Linear mixed-effects regressions with repeated measures were used to assess the evolution of spinal and cortical activations with increasing force levels. We found a significant increase (*P*<0.001) in ipsilateral spinal cord activation (estimated means: low 28.3, medium 57.8, high 78.1, Fig. 2A), post-hoc tests confirmed a significant difference between all three force levels (Tab. 1). Similarly, also contralateral M1 exerted a force level dependent increase (low 215, medium 246, high 286) in focal brain activation (*P*<0.001) and post-hoc tests also confirmed a significant difference between all three force levels (Tab.1, Fig. 2B). For SMA (low 130, medium 128, high 139) the model also revealed a significant dependence on the force levels (*P*=0.019), but the post-hoc tests revealed only a significant difference between medium and high (*P*=0.027, Tab. 1). For PMV (low 106, medium 108, high 110), there were no significant differences between the force levels (*P*=0.61, Fig. 2B, Tab. 1).

**Figure 2.**
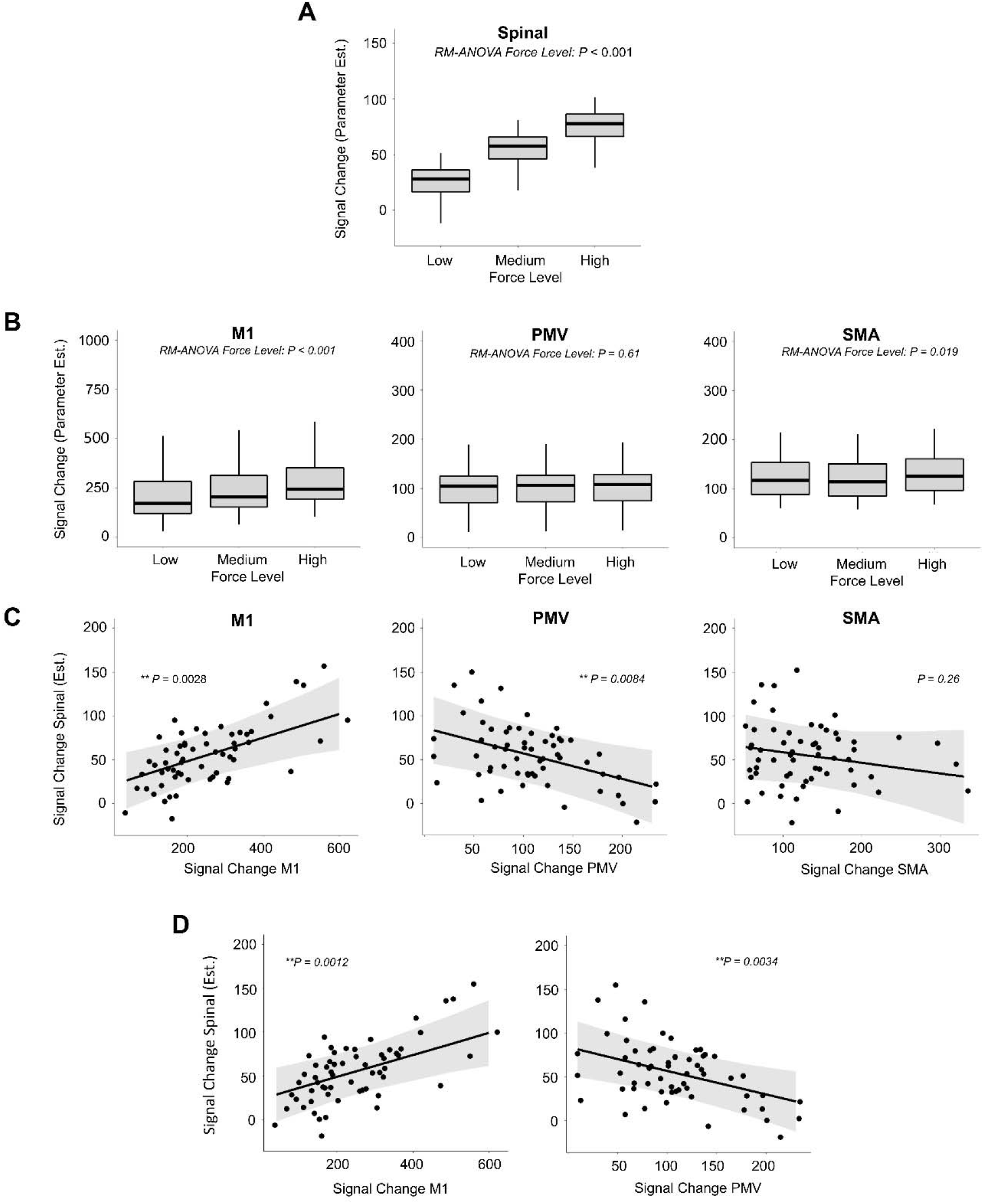
Spinal cord and cortical brain activation and their association during force generation. Linear mixed model repeated measures analyses. **A** Estimated spinal activation depending on the three force levels. **B** Estimated cerebral activation in the three cerebral regions M1, PMV, SMA depending on the three force levels **C** Effect plots showing association between cerebral activation in M1, PMV and SMA and the estimated spinal activation. Shown are the results of individual linear mixed-effects models in which M1, PMV or SMA activation was separately related to the spinal activation, respectively (univariate). **D** Effect plots of the combined analysis of M1 and PMV activation contributing to spinal cord activation (multivariate). Importantly, M1 and PMV activations were not correlated. *P*-values are given for the factors of interest. \**P*<0.05, \*\**P*<0.01.

**Table 1.** Pairwise comparisons of BOLD activation between force levels. P values are given uncorrected for pairwise post-hoc comparisons between force levels for spinal, M1, SMA, PMV activations.

| Comparisons | Spinal | M1 | SMA | PMV |
| --- | --- | --- | --- | --- |
| low - medium | 0.0001** | 0.0001** | 0.80 | 0.93 |
| medium - high | 0.0051** | <0.0001*** | 0.027* | 0.82 |
| low - high | <0.0001*** | <0.0001*** | 0.11 | 0.60 |

### Relationships between spinal cord and cortical brain activation

Finally, linear mixed-effects regressions with repeated measures were used to address the relationship between cerebral activation in M1, PMV and SMA and spinal cord activation. We found a significant and positive correlation between M1 activation and spinal cord activation (P=0.0028, Fig. 2C). In contrast, PMV was negatively related to spinal cord activation (P=0.0084, Fig. 2C). SMA did not exhibit any significant association with the amount of spinal activation (P=0.26, Fig. 2C). To further analyze the influence of M1 and PMV activation on spinal activation, a linear mixed model with M1 and PMV BOLD responses as fixed factors was estimated. This model confirmed the negative association of PMV activation to spinal activation (P=0.0034) independent from the influence of M1 (P=0.0012). A potential interaction of M1 and PMV onto spinal activation was not evident (P=0.13). Also, M1 and PMV did not show any cross-correlation (P=0.60).

## Discussion

The most remarkable novel finding of the present study is the significant influence of PMV on spinal cord activation during force generation. This involvement was independent from the positive relationship between activation in the contralateral primary motor cortex (M1) and the ipsilateral spinal cord. More precisely, PMV activation was negatively linked with spinal cord activation. PMV and M1 activations were not correlated with each other. These findings suggest that the PMV might be connected to lower cervical spinal cord motoneurons in segments contributing to distal upper limb functions, most likely via indirect, poly-synaptic pathways. The supplementary motor area (SMA), an alternative secondary motor region, did not exhibit comparable associations with spinal cord activation during force generation.

The presented force generation task activated a bilateral and widespread cerebral network including contralateral sensorimotor cortices, and premotor areas such as the PMV, SMA, dorsolateral prefrontal cortices and areas of the posterior parietal lobe along the intraparietal sulcus. These areas are all well in line with previous imaging data, derived from power grip ^5,7,57^ and precision grip force tasks ^11,12^. Compared to this cortical brain network, a systematic analysis of cortico-spinal interactions during force generation was not available so far. The present study aimed at answering two main questions: Does activation in the contralateral M1 show a significant association with ipsilateral spinal cord activation? Might premotor cortices also show an impact on spinal cord activation during force generation?

Both M1 and the spinal cord showed a force level dependent increase in the spatial extent of activated voxels and in the strength of activation. Force level dependent dynamics in M1 activation are well in line with previous animal data ^14,58^ and data from human power grip force experiments ^3,5,7,57,59^. On the spinal level, similar patterns were observed at increasing force levels: Not only the peak activation increased with higher force levels, but also the extent of activated spinal tissue showed gradually increases suggesting that more and more motor units and muscles got active to generate the target force ^60–62^. In fact, several muscle groups work in synergy during finger flexion for power grip. The primary finger flexors at lower force levels are the flexor digitorum superficialis and the flexor digitorum profundus muscle. They are innervated by motor neurons residing in C6 to T1 spinal cord segments. The palmaris longus muscle and the lumbrical muscles, innervated by C7 to T1 spinal cord segments, can contribute. Hence, finger flexion is primarily located in spinal cord segments C6-T1 ^63–65^. The observed BOLD signals, mainly located in the C6 to C8 spinal cord segments (corresponding to the C5–C7 vertebral level), agree with these anatomical considerations. When comparing the evolution of M1 and spinal activation under increasing forces, the numerical increase in activation was more pronounced at the spinal level when compared to M1 suggesting that the spinal activation is more directly linked to the applied force than M1 activity.

In addition to the force level dependent dynamics in M1 and the spinal cord, we found a significant positive correlation between M1 activation and spinal cord activation. This matches widely established concepts of M1 as the main origin of the CST. Similar results of such cortico-spinal co-activation in functional MRI have been recently obtained during complex finger movements ^26^.

In addition to M1, force generation also involved bilateral PMV and SMA. However, in contrast to M1, PMV did not show any significant modulation while target forces increased. For SMA, post-hoc tests did only show marginal effects between medium and high force level. Stable activations in the dorsal premotor cortex were not detected at all. Similar results have already been reported by one previous study ^5^. Therefore, the subsequent analyses for cortico-spinal coupling between premotor regions and the spinal cord were limited to PMV and SMA.

We observed a significant negative association between PMV activation and spinal activation at lower C7 to upper C8 spinal cord segments. This finding is novel in two important aspects.

First, it provides first empirical functional data in humans to support the view that PMV might influence spinal motor neuronal activity at lower cervical segments which are involved in distal upper limb functions. Animal tracing data in non-human primates have reported PMV analogues to primarily terminate in the upper segments of the spinal cord only ^38,39,48,49^. Poly-synaptic connections have been proposed to reach distal forelimb segments along different inter-neural indirect routes ^44–47^. Human data on the contribution of premotor cortices such as PMV to spinal processing are still limited. For instance, using TMS, peripheral nerve stimulation and monosynaptic reflex analysis, one study has addressed the input convergence between peripheral afferents and cortico-spinal inputs originating from PMV onto proprio-spinal neurons. This study has suggested the existence of descending influences onto the proprio-spinal system not only for M1 but also for PMV ^52^. Tractography data have recently argued – despite critical limitations in the spatial resolution – for the existence of direct connections between premotor areas including PMV and SMA and even lower cervical segments ^53^.

Second, the present results with correlated PMV activity and spinal activity and uncorrelated PMV and M1 activities significantly extend previously published network data during force generation. One study has used effective connectivity analyses to characterize information flow between M1 and premotor cortices. Premotor-M1 coupling has been found to increase linearly from lower to higher grip forces. It has been speculated that premotor cortices might be involved in force generation while primarily modulating the output of M1 to the spinal motor neurons ^59^. Another study has discussed widespread human cortical and subcortical brain networks involved in force generation. The actual data indicate now that the PMV might not only contribute to networks at the cortical level, but it may also exert additional, more direct influences onto the spinal cord. Hence, brain activity involved in force generation would converge towards the spinal cord not only via M1 and its CST as the main outflow tract, but cortico-fugal information outflow might be also mediated via premotor cortices. This would further strengthen the *CST bypass* concepts which have emerged over the last years in stroke recovery research. Studies have consistently evidenced that the integrity of secondary CSTs originating from premotor cortices might influence residual motor output and recovery after stroke ^66^.

How can we interpret the negative correlation between PMV and spinal cord activation? Data in non-human primates have suggested the existence of cortico-spinal connections targeting the upper cervical spinal cord. Results are variable regarding the precise topography, potential relay nodes and involved secondary inter-neuronal networks ^67^. For instance, non-human primate concepts have been developed proposing a poly-synaptic course via segmental interneurons, proprio-spinal neurons, reticulo-spinal and rubro-spinal neurons ^44^. Evidence suggesting proprio-spinal inter-neurons as important inhibitory target networks for top-down PMV influence in humans comes from a recent spatial facilitation study combining TMS and peripheral nerve stimulation on the upper limb ^52^. Finally, the absence of retrograde cortical degeneration in PMV in patients after spinal cord injury and absent motor responses after PMV electrical stimulation have argued against direct mono-synaptic connections between PMV and the spinal cord ^53^. Therefore, we speculate that poly-synaptic trajectories originating from PMV with long-range excitatory pathways converging onto inhibitory spinal networks which modulate spinal motor neurons, are most likely to transmit the cortico-spinal influence from PMV during force generation.

There are several limitations to note. First, they include possible artifacts in the spinal images related to movement and physiological signals ^68^. The analysis pipeline was adapted according to established protocols for spinal fMRI artifact minimization. ^23,24,69–71^. Second, smoothing of the fMRI data might complicate the precise in-plane localization and slight shifts in the single-subject activation maps during normalization might explain why activation peaks at group level are distributed between the anterior and posterior horn. Third, the cerebral and spinal field-of-view was limited due to technical restrictions. Therefore, it was not possible to investigate the influence of other brain regions, e.g., basal ganglia or the cerebellum on the spinal activation.

## Materials and Methods

### Subjects

21 young volunteers participated in the study. 1 subject was excluded from further analysis because of an accidental MRI finding. 20 young volunteers were included into the analysis. All subjects were right-handed according to the Edinburgh handedness inventory ^72^, had normal or corrected to normal vision and reported no neurological or musculoskeletal diseases or contraindications to MRI. The study was conducted following the Declaration of Helsinki and approved by the local ethics committee of the Medical Association of Hamburg (PV6026). All subjects gave written informed consent and received monetary compensation.

### Motor Task

We employed an fMRI block design with three experimental conditions (block length 15 s) and interleaved resting baselines (Fig. 3A) using Psychtoolbox version 3.0.16, ran in Octave version 4.0.3. In the experimental conditions, the subjects were asked to perform visual cued whole hand grips with their right hand with three different predefined force levels low, medium and high corresponding to 30%, 50%, 70% of the maximum output measurement (linear force measurement) covered by an MRI compatible grip force response device (Grip Force Bimanual, Current Design, Inc, Philadelphia, PA). All three experimental conditions had the same time course. First, the subjects were informed via a video screen, visible through a mirror attached to the head coil of the MR scanner, which force level they had to reach in the upcoming activation block. After 1.5 s, the instruction text was replaced by an empty circle. After a variable delay of 1.5, 2.0, or 2.5 s, a white cross blinked under the empty circle at 0.8Hz (the cross appeared for 0.625 s). The subjects then performed almost isometric hand grips with the right hand synchronized with the white cross. When they reached 90% of the correct force level, the empty white circle changed to a solid white circle. The instruction was to maintain this strength for the duration of the appearance of the white cross. After 15 s, the blinking cross and circle were replaced by a black screen, which told the subjects to stop the hand movements and rest until the next instruction appeared.

**Figure 3.**
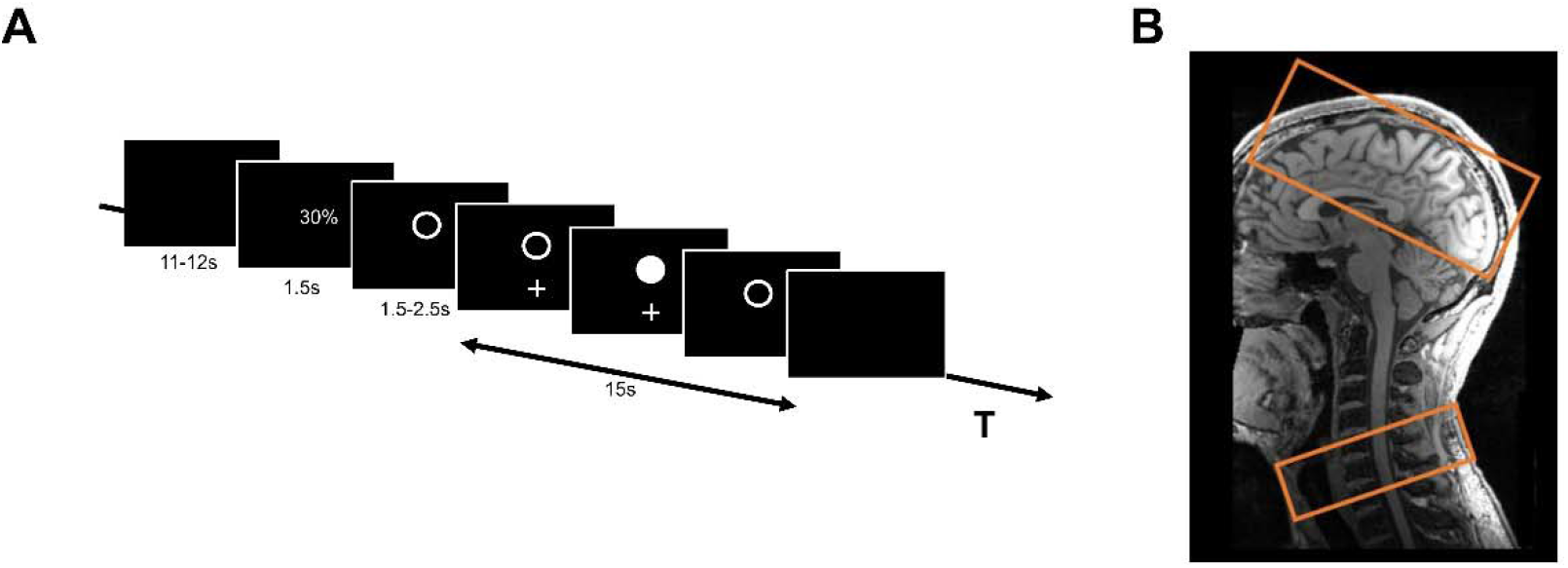
Schematic representation. **A** Task design; **B** Exemplary representation of the position of the two sub-volumes

The duration of this resting (= baseline) condition depended on a variable delay between the end of the instruction text and the start of the cue lasting between 11 and 12 s, resulting in an inter-block interval of 15 s. The sequence of the three different force levels was pseudorandomized. Each fMRI session consisted of 490 images preceded by five dummy images allowing the MR scanner to reach a steady state in T2* contrast. After acquiring the dummy images, the experiment started with a baseline condition. The whole session consisted of 36 blocks (12 for each force level, lasting 15 s) and 36 baseline blocks + instruction conditions (lasting 15 s). The session was repeated twice. Each session lasted approximately 18.4 minutes. The subjects were trained outside the scanner to familiarize themselves with the task. During this training session, they were trained to perform the hand movements at the defined frequency and reach the correct force levels without overshooting. After the training session, the subjects received feedback on their performance to achieve a performance improvement. Inside the MR scanner, the subject’s hand and arm were placed in a comfortable position on the belly of the subject. If necessary, medical tape fixed the force response device on the right hand. During the whole session, the force production was recorded for later analysis.

### MRI data acquisition

A 3T Prisma MRI scanner (Siemens Healthineers, Erlangen, Germany) and a 64-channel combined head-neck coil were used to acquire cerebral and spinal imaging data. The magnet’s iso-center was set on the lower part of the chin of the subject, approximately centered to vertebral C2/C3, but in some cases, it had to be further adjusted depending on the subject’s height. The imaging modalities included high-resolution T1-weighted, T2*-weighted, and task-evoked fMRI images. For the T1-weighted sequence, a 3-dimensional magnetization-prepared rapid gradient echo (3D-MPRAGE) sequence was used, which covered the head and neck (cervical spine and upper part of the thoracic spine) with the following parameters: repetition time (TR)=2300ms, echo time (TE)=3.4ms, flip angle 9°, 236 coronal slices, 320 axial slices, with a voxel size of 1.0×1.0×1.0 mm^3^. The T2*-weighted image (MEDIC sequence) covered the lower part of the cervical spine, centered on the cervical vertebra C6, with the following parameters: TR=307ms, TE=21ms, flip angle 20°, eight axial slices, with a voxel size of 0.5×0.5×5.0 mm^3^, the slices were positioned identical to the spinal slices of the functional acquisitions, see below. For fMRI, a combined cortico-spinal fMRI protocol based on echo-planar imaging (EPI) was used to record BOLD responses in the brain and spinal cord ^21,73^, 32 slices, divided into two sub-volumes (Fig. 3B), were acquired. These two sub-volumes have different geometry, timing parameter ^21,74^ and shim settings. The shim settings were determined using a field map acquisition and a dedicated shim algorithm ^74,75^. The upper volume included 24 axial slices (voxel-size: 2.0×2.0×2.0 mm3, 1 mm gap between slices) in the brain, which was initially oriented along the anterior-posterior commissure axis and, if necessary, tilted to cover the primary and secondary motor areas and if possible, the visual cortex. The lower sub-volume consisted of 8 axial slices (voxel-size: 1.0×1.0×5.0 mm3, no gap between slices), centered at the vertebral body of C6 and covered the vertebral bodies of C5, C6, and C7. The whole sequence was measured with the following parameters: TR=2231ms, TE=30ms (brain) and 31ms (spinal), flip angle=75°. Additionally, we measured one whole-brain EPI volume with the following parameters: TR=2385ms, TE=30ms, flip angle 75°, and 36 axial slices with a voxel size of 2.0×2.0×2.0 mm3 with 1 mm gap between slices. During the fMRI sessions, pulse, ECG, respiration, and the trigger signal were recorded (sampling rate=400Hz) using the Physlog-function (Ideacmdtool, provided by Siemens Healthineers, Erlangen, Germany) and respiratory, ECG, and pulse measurement devices provided by Siemens Healthineers, Erlangen, Germany)

### Behavioral data

The individual maximum grip force (whole hand grip) of the right hand was measured with a JAMAR Hand Dynamometer (built by Patterson Medical, Warrenville, USA).

### Image preprocessing

Brain and spinal cord images were pre-processed separately. The brain fMRI images were pre-processed using the Oxford Center for fMRI of the Brain’s (FMRIB) Software Library (FSL) version 6.0.4 ^76^. The whole-brain EPI-image of each subject was linear co-registered to the brain-extracted high-resolution T1-image of each subject, and the individual T1-image was linear co-registered to the MNI152-T1-2mm image provided by the FSL library. The transformation matrices were concatenated for further preprocessing steps. The first 5 dummy volumes of the task-related fMRI images were discarded. The mean fMRI-image was registered to the whole brain EPI, and then the concatenated transformation matrices were used for registration on the MNI152-T1-2mm image. The fMRI images were further pre-processed with motion correction using MCFLIRT ^77^, and the images from both sessions were concatenated into one time series at the subject level.

The spinal fMRI images were pre-processed using the Spinal Cord Toolbox, version 5.2 ^24^ and FSL version 6.0.4 ^76^. The spinal fMRI images were cropped with the spinal cord at the center of the image. Motion correction was performed using 2 phases of movement correction. MCFLIRT ^77^ was used for the first phase of motion correction with spline interpolation and a normalized correlation cost function. The images across the two runs were realigned to the first image of the first run with a three-dimensional rigid-body realignment. To correct for slice-independent motion due to the non-rigidity of the cervical spine and physiological motion from swallowing and the respiratory cycle, the second phase of motion correction was performed with two-dimensional rigid realignment independently for each axial slice ^71,78^. The images from both sessions were concatenated into one time series at the subject level. The spatial normalization from native to standard space was performed using tools from the open-source Spinal Cord Toolbox ^24,78^: The C5 and C7 vertebrae in the structural T2* images of the cervical spine were manually identified, and the spinal cord was automatically identified and segmented. The structural images were then normalized to the PAM50_T2s-template (resolution=0.5×0.5×0.5 mm^3^) ^78,79^. After motion correction, the mean functional image was segmented to identify the spinal cord. The resulting binary spinal cord mask and the reversed deformation fields of the structural normalization were used to register the PAM50_T2-template on the mean functional image. The inverted resulting deformation field was then used to normalize the functional images and other images (e.g., cope-images) to PAM50-space. The normalized images were visually inspected for quality control at each step.

### Physiological noise modeling

Cardiac and respiratory cycles are significant noise sources in spinal cord fMRI and can confound signal detection ^78^. To account for this noise, cardiac signals (pulse), respiratory signals, and MRI triggers were collected during scanning. The SPM (SPM12) based PhysIO Toolbox version 8.0.1 ^70^, ran in MATLAB version R2018a was used to calculate the noise regressors. This toolbox uses a model-based physiological noise correction, which uses retrospective image correction (RETROICOR) of physiological motion effects ^80^, heart rate variability ^81^, and respiratory volume per time ^82^. Based on the physiological signals, 18 noise regressors were generated. A cerebrospinal fluid (CSF) regressor was also generated from the CSF signal surrounding the spinal cord using a subject-specific CSF mask generated from the PAM50_csf-template ^78,79^.

### Data analysis: First and second-level analyses

Two different first- and second-level analyses were performed. The analyses of the cerebral and spinal images were conducted separately. For both analyses, the same explanatory variables (EVs) were used in the design matrices of the general linear model (GLM) analyses. For the first first-level analysis, the recorded force production, which was recorded during the MRI sessions, was further analyzed. For each volume (TR=2.231s), the mean of the maximum force produced was calculated and used as EV in the design matrix and hereinafter referred to as mean EV. In the second first-level analysis, the force levels low, medium, and high were used as EVs. In both analyses, the temporally jittered instruction period was separately modeled as an additional EV but not further analyzed in the group analysis ^83^. For the group analysis, separate analyses for the brain and spinal cord data were performed.

For the brain images, the motion-corrected functional images were spatially smoothed with a Gaussian kernel of 5 mm full width half maximum (FWHM) and high-pass filtered (90 s) using the fMRI Expert Analysis TOOL (FEAT v6.00) ^84,85^. Statistical maps of the pre-processed time series were generated using FMRIB’s improved Linear Model (FILM) with pre-whitening ^78,85^. The design matrices included the hemodynamic response function (gamma convolution, phase 0 s, standard deviation 3 s, mean lag 6 s) convolved task vectors as EVs, motion parameters, and motion outliers, determined using *fsl_motion_outliers*, were included as covariates of no interest. For the second-level group analysis, spatial normalization of the statistical images from the subject-level analyses to the MNI template was performed. Group average activation maps for each contrast were generated with the demeaned individual grip force of the right hand as an additional covariate using FMRIB’s Local Analysis of Mixed-effects (FLAME) stages 1 and 2 ^84,86^. The group average activation maps were thresholded using a Z-Score>3.5 with a cluster significance threshold of *P*<0.05 to correct for multiple comparisons, in line with previous recommendations ^87^.

Additionally, three binary masks were generated from the HMAT-Template (Human motor area template) ^88^ covering left primary motor cortex (M1), left ventral premotor cortex (PMV) and left supplementary motor area (SMA). The masks were used to detect the peak voxels in the individual subject-specific activation maps for the mean EV. Each peak voxel location was manually checked for plausibility and, if necessary, manually corrected. The mean parameter estimates of the three different force levels were extracted using spheres of a radius of 5 mm centered on the peak coordinates.

For the spinal images, the motion-corrected functional images were spatially smoothed with a Gaussian kernel of 2×2×5 mm FHWM and the spinal cord was extracted from the data using a spinal cord mask, which was created from the PAM50_cord_template and spatial transformed in the subject-specific space. The data were further analyzed with FEAT from the FMRIB software library ^85^ and were high-pass filtered (90 s). The statistical maps of the pre-processed time series were generated using FILM with pre-whitening ^78,85^. The design matrices included the hemodynamic response function (gamma convolution, phase 0 s, standard deviation 3 s, mean lag 6 s), convolved task vectors as EVs, the physiological noise regressors, the CSF time series, and the motion parameters as covariates of no interest. For the second-level group analysis, spatial normalization of the statistical images from the subject-level analyses to the PAM50-template was performed. Group average activation maps for each contrast were generated with the demeaned individual maximum force as an additional covariate using FLAME stages 1 and 2 ^78,86^. The group average activation maps were thresholded using a Z-Score>2.5 (lower Z-threshold than in the cerebral images, adapted to the lower detected activation in the spinal images, such as in ^78^) with a cluster significance threshold of *P*<0.05 to correct for multiple comparisons. Additionally, a binary mask was generated to analyze the individual activation maps, which covered the right hemi-cord between the vertebral-level C5 and C7. The mask was used to detect the peak voxel in the individual subject-specific activation maps for the mean EV. The mean parameter estimates of the three different force levels were extracted using in-plane spheres with a radius of 1.5 mm centered on the peak coordinates.

### Further statistical analysis

The statistical package R 4.0.4 ^89^ was used for further statistical analysis. To analyze brain (M1, PMV, SMA) or spinal cord activation under varying target forces, linear mixed-effects models with repeated measures (package lme4) were used, with brain activation values as the dependent variable, force levels as the independent factor of interest and subject as a random factor. Post-hoc tests with pairwise comparisons between force levels were carried out using emmeans. To analyze the relationship between cerebral and spinal activation, the following linear mixed-effects models were fitted with spinal activation as the dependent variable and activation values as independent, fixed factor of interest and subject and force levels as random factors. 1) Three separate models were computed to analyze the relationship between spinal activation and regional brain activation of the three regions of interest (M1, SMA, PMV). 2) One model was additionally estimated combining significant cerebral activations from the univariate models. 3) The combined model was further explored by adding an interaction term for the relevant brain activations. Statistical significance was assumed at P<0.05. Leave-one-out model analyses of the fitted models were applied to ensure robustness of the findings with statistical significance remaining stable when iteratively excluding all single subjects.

## Author contributions

HB and CG designed the study. JFE and HB designed and developed the motor task. HB performed all MRI experiments and data acquisition. JFI and YC developed the fMRI sequences and implemented them on the MRI. HB and RS performed data analysis and statistical analysis. HB, RS, AT, CB and CG performed data interpretation. HB and RS wrote the first version of the manuscript.

## Data and materials availability

All data needed to evaluate the conclusions in the paper are present in the paper and/or the Supplementary Materials. Processed brain and spinal cord activation data and demographic data to reproduce the findings are available from the corresponding author on reasonable request.

## References

1. Ehrsson, H. H. et al. Cortical activity in precision-versus power-grip tasks: An fMRI study. J. Neurophysiol. 83, 528–536 (2000).

2. Spraker, M. B., Corcos, D. M. & Vaillancourt, D. E. Cortical and subcortical mechanisms for precisely controlled force generation and force relaxation. Cereb. Cortex 19, 2640–2650 (2009).

3. Cramer, S. C. et al. Motor cortex activation is related to force of squeezing. Hum. Brain Mapp. 16, 197–205 (2002).

4. Mima, T., Simpkins, N., Oluwatimilehin, T. & Hallett, M. Force level modulates human cortical oscillatory activities. Neurosci. Lett. 275, 77–80 (1999).

5. Keisker, B., Hepp-Reymond, M. C., Blickenstorfer, A., Meyer, M. & Kollias, S. S. Differential force scaling of fine-graded power grip force in the sensorimotor network. Hum. Brain Mapp. 30, 2453–2465 (2009).

6. Keisker, B., Hepp-Reymond, M.-C. C., Blickenstorfer, A. & Kollias, S. S. Differential representation of dynamic and static power grip force in the sensorimotor network. Eur. J. Neurosci. 31, 1483–1491 (2010).

7. Dai, T. H., Liu, J. Z., Saghal, V., Brown, R. W. & Yue, G. H. Relationship between muscle output and functional MRI-measured brain activation. Exp. Brain Res. 140, 290–300 (2001).

8. Nowak, D. A., Hermsdörfer, J., Marquardt, C. & Fuchs, H. H. Grip and load force coupling during discrete vertical arm movements with a grasped object in cerebellar atrophy. Exp. Brain Res. 145, 28–39 (2002).

9. Nowak, D. A., Hermsdörfer, J. & Topka, H. Deficits of predictive grip force control during object manipulation in acute stroke. J. Neurol. 250, 850–860 (2003).

10. Duque, J. et al. Correlation between impaired dexterity and corticospinal tract dysgenesis in congenital hemiplegia. Brain 126, 732–747 (2003).

11. Olivier, E., Davare, M., Andres, M. & Fadiga, L. Precision grasping in humans: from motor control to cognition. Current Opinion in Neurobiology vol. 17 644–648 (2007).

12. Castiello, U. & Begliomini, C. The cortical control of visually guided grasping. Neuroscientist vol. 14 157–170 (2008).

13. Evarts, E. V., Fromm, C., Kroller, J. & Jennings, V. A. A. Motor cortex control of finely graded forces. J. Neurophysiol. 49, 1199–1215 (1983).

14. Evarts, E. V. Relation of pyramidal tract activity to force exerted during voluntary movement. J. Neurophysiol. 31, 14–27 (1968).

15. Hepp-Reymond, M. C., Kirkpatrick-Tanner, M., Gabernet, L., Qi, H. X. & Weber, B. Context-dependent force coding in motor and promotor cortical areas. in Experimental Brain Research vol. 128 123–133 (1999).

16. Dafotakis, M., Sparing, R., Eickhoff, S. B., Fink, G. R. & Nowak, D. A. On the role of the ventral premotor cortex and anterior intraparietal area for predictive and reactive scaling of grip force. Brain Res. 1228, 73–80 (2008).

17. Davare, M., Andres, M., Clerget, E., Thonnard, J. L. & Olivier, E. Temporal dissociation between hand shaping and grip force scaling in the anterior intraparietal area. J. Neurosci. 27, 3974–3980 (2007).

18. Spraker, M. B., Yu, H., Corcos, D. M. & Vaillancourt, D. E. Role of individual basal ganglia nuclei in force amplitude generation. J. Neurophysiol. 98, 821–834 (2007).

19. Spraker, M. B. et al. Specific cerebellar regions are related to force amplitude and rate of force development. Neuroimage 59, 1647–1656 (2012).

20. Finsterbusch, J., Eippert, F. & Büchel, C. Single, slice-specific z-shim gradient pulses improve t2*-weighted imaging of the spinal cord. Neuroimage 59, 2307–2315 (2012).

21. Finsterbusch, J., Sprenger, C. & Büchel, C. Combined T2*-weighted measurements of the human brain and cervical spinal cord with a dynamic shim update. Neuroimage 79, 153–161 (2013).

22. Islam, H., Law, C. S. W., Weber, K. A., Mackey, S. C. & Glover, G. H. Dynamic per slice Shimming for Simultaneous Brain and Spinal Cord fMRI. Magn. Reson. Med. 81, 825–838 (2019).

23. Eippert, F., Kong, Y., Jenkinson, M., Tracey, I. & Brooks, J. C. W. Denoising spinal cord fMRI data: Approaches to acquisition and analysis. Neuroimage 154, 255–266 (2017).

24. De Leener, B. et al. SCT: Spinal Cord Toolbox, an open-source software for processing spinal cord MRI data. Neuroimage 145, 24–43 (2017).

25. Vahdat, S. et al. Resting-state brain and spinal cord networks in humans are functionally integrated. PLoS Biol. 18, (2020).

26. Takasawa, E., Abe, M., Chikuda, H. & Hanakawa, T. A computational model based on corticospinal functional MRI revealed asymmetrically organized motor corticospinal networks in humans. Commun. Biol. 5, 664 (2022).

27. Landelle, C. et al. Investigating the Human Spinal Sensorimotor Pathways Through Functional Magnetic Resonance Imaging. Neuroimage 245, 118684 (2021).

28. Mima, T. & Hallett, M. Corticomuscular coherence: A review. Journal of Clinical Neurophysiology vol. 16 501–511 (1999).

29. Gerloff, C. et al. Coherent Corticomuscular Oscillations Originate From Primary Motor Cortex: Evidence from Patients with Early Brain Lesions. Hum. Brain Mapp. 27, 789–798 (2006).

30. Rossiter, H. E. et al. Changes in the location of cortico-muscular coherence following stroke. NeuroImage Clin. 2, 50–55 (2013).

31. Belardinelli, P., Laer, L., Ortiz, E., Braun, C. & Gharabaghi, A. Plasticity of premotor cortico-muscular coherence in severely impaired stroke patients with hand paralysis. NeuroImage Clin. 14, 726–733 (2017).

32. Lemon, R. N. The Cortical “Upper Motoneuron” in Health and Disease. Brain Sci. 11, 619 (2021).

33. Lemon, R. N. Descending Pathways in Motor Control. Annu. Rev. Neurosci. 31, 195–218 (2008).

34. Catsman-Berrevoets, C. E. & Kuypers, H. G. J. M. Cells of origin of cortical projections to dorsal column nuclei, spinal cord and bulbar medial reticular formation in the rhesus monkey. Neurosci. Lett. 3, 245–252 (1976).

35. Biber, M. P., Kneisley, L. W. & LaVail, J. H. Cortical neurons projecting to the cervical and lumbar enlargements of the spinal cord in young and adult rhesus monkeys. Exp. Neurol. 59, 492–508 (1978).

36. Murray, E. A. & Coulter, J. D. Organization of corticospinal neurons in the monkey. J. Comp. Neurol. 195, 339–365 (1981).

37. Dum, R. P. & Strick, P. L. The origin of corticospinal projections from the premotor areas in the frontal lobe. J. Neurosci. 11, 667–89 (1991).

38. He, S.-Q. Q., Dum, R. P. & Strick, P. L. Topographic Organization of Corticospinal Projections from the Frontal Lobe□: Motor Areas on the Lateral Surface of the Hemisphere. J. Neurosci. 13, 952–80 (1993).

39. He, S. Q., Dum, R. P. & Strick, P. L. Topographic organization of corticospinal projections from the frontal lobe: Motor areas on the medial surface of the hemisphere. J. Neurosci. 15, 3284–3306 (1995).

40. Dum, R. P. & Strick, P. L. Spinal cord terminations of the medial wall motor areas in macaque monkeys. J. Neurosci. 16, 6513–6525 (1996).

41. Maier, M. A. et al. Differences in the corticospinal projection from primary motor cortex and supplementary motor area to macaque upper limb motoneurons: An anatomical and electrophysiological study. Cereb. Cortex 12, 281–296 (2002).

42. Innocenti, G. M. et al. Diversity of Cortico-descending Projections: Histological and Diffusion MRI Characterization in the Monkey. Cereb. Cortex 29, 788–801 (2019).

43. Borra, E., Belmalih, A., Gerbella, M., Rozzi, S. & Luppino, G. Projections of the hand field of the macaque ventral premotor area F5 to the brainstem and spinal cord. J. Comp. Neurol. 518, 2570–2591 (2010).

44. Isa, T., Kinoshita, M. & Nishimura, Y. Role of direct vs. indirect pathways from the motor cortex to spinal motoneurons in the control of hand dexterity. Front. Neurol. 4, 1–9 (2013).

45. Isa, T., Ohki, Y., Seki, K. & Alstermark, B. Properties of propriospinal neurons in the C3-C4 segments mediating disynaptic pyramidal excitation to forelimb motoneurons in the macaque monkey. J. Neurophysiol. 95, 3674–3685 (2006).

46. Isa, T., Ohki, Y., Alstermark, B., Pettersson, L.-G. G. & Sasaki, S. Direct and indirect cortico-motoneuronal pathways and control of hand/arm movements. Physiology 22, 145–153 (2007).

47. Alstermark, B., Lundberg, A. & Sasaki, S. Integration in descending motor pathways controlling the forelimb in the cat - 11. Inhibitory pathways from higher motor centres and forelimb afferents to C3-C4 propriospinal neurones. Exp. Brain Res. 56, 293–307 (1984).

48. Wise, S. P. The ventral premotor cortex, corticospinal region C, and the origin of primates. Cortex 42, 521–524 (2006).

49. Morecraft, R. J. et al. Terminal organization of the corticospinal projection from the lateral premotor cortex to the cervical enlargement (C5–T1) in rhesus monkey. J. Comp. Neurol. 527, 2761–2789 (2019).

50. Fleischmann, R., Triller, P., Brandt, S. A. & Schmidt, S. H. Human Premotor Corticospinal Projections Are Engaged in Motor Preparation at Discrete Time Intervals: A TMS-Induced Virtual Lesion Study. Front. Neuroergonomics 2, (2021).

51. Teitti, S. et al. Non-primary motor areas in the human frontal lobe are connected directly to hand muscles. Neuroimage 40, 1243–1250 (2008).

52. Giboin, L. S. et al. Corticospinal control from M1 and PMv areas on inhibitory cervical propriospinal neurons in humans. Physiol. Rep. 5, e13387 (2017).

53. Usuda, N. et al. Quantitative comparison of corticospinal tracts arising from different cortical areas in humans. Neurosci. Res. 183, 30–49 (2022).

54. Ito, K. L. et al. Corticospinal Tract Lesion Load Originating From Both Ventral Premotor and Primary Motor Cortices Are Associated With Post-stroke Motor Severity. Neurorehabil. Neural Repair 36, 179–182 (2022).

55. Schulz, R. et al. Assessing the Integrity of Corticospinal Pathways From Primary and Secondary Cortical Motor Areas After Stroke. Stroke 43, 2248–2251 (2012).

56. Newton, J. M. et al. Non-invasive mapping of corticofugal fibres from multiple motor areas-relevance to stroke recovery. Brain 129, 1844–1858 (2006).

57. Alahmadi, A. A. S. et al. Blood Oxygenation Level-Dependent Response to Multiple Grip Forces in Multiple Sclerosis: Going Beyond the Main Effect of Movement in Brodmann Area 4a and 4p. Front. Cell. Neurosci. 15, (2021).

58. Hepp-Reymond, M. C., Wannier, T. M. J., Maier, M. A. & Rufener, E. A. Chapter 37 Sensorimotor cortical control of isometric force in the monkey. in Progress in Brain Research (eds. Allum, J. H. J. & Hulliger, M.) vol. 80 451–463 (Elsevier, 1989).

59. Saleh, S., Jiang, Z. & Yue, G. H. Motor Control Network Effective Connectivity in Regulating Muscle Force Output. Nat. Sci. 13, 9–17 (2021).

60. Purves, D. et al. The Regulation of Muscle Force. in Neuroscience, 2nd edition (ed. Sunderland) (Sinauer Associates, 2001).

61. Heckman, C. J. & Enoka, R. M. Chapter 6 Physiology of the motor neuron and the motor unit. in Handbook of Clinical Neurophysiology (ed. Eisen, A.) vol. 4 119–147 (Elsevier, 2004).

62. Mendell, L. M. The size principle: A rule describing the recruitment of motoneurons. J. Neurophysiol. 93, 3024–3026 (2005).

63. Ghez, C. & Krakauer, J. Back 33 The Organization of Movement. in Principles of neural science 653–673 (McGraw-Hill, 2000).

64. Moore, K. L., Dalley, A. F. & Agur, A. M. R. Clinically Oriented Anatomy Seventh Edition. (Wolters Kluwer, 2014).

65. Stifani, N. Motor neurons and the generation of spinal motor neuron diversity. Front. Cell. Neurosci. 8, 1–22 (2014).

66. Koch, P., Schulz, R. & Hummel, F. C. Structural connectivity analyses in motor recovery research after stroke. Ann. Clin. Transl. Neurol. 3, 233–244 (2016).

67. Zholudeva, L. V. et al. Spinal Interneurons as Gatekeepers to Neuroplasticity after Injury or Disease. J. Neurosci. 41, 845–854 (2021).

68. Fratini, M., Moraschi, M., Maraviglia, B. & Giove, F. On the impact of physiological noise in spinal cord functional MRI. J. Magn. Reson. Imaging 40, 770–777 (2014).

69. Brooks, J. C. W. et al. Physiological noise modelling for spinal functional magnetic resonance imaging studies. Neuroimage 39, 680–692 (2008).

70. Kasper, L. et al. The PhysIO Toolbox for Modeling Physiological Noise in fMRI Data. J. Neurosci. Methods 276, 56–72 (2017).

71. Cohen-Adad, J., Rossignol, S. & Hoge, R. Slice-by-slice motion correction in spinal cord fMRI: SliceCorr. Proc. 17th Sci. Meet. Int. Soc. Magn. Reson. Med. Honolulu 3181 (2009).

72. Oldfield, R. C. The assessment and analysis of handedness: The Edinburgh inventory. Neuropsychologia 9, 97–113 (1971).

73. Tinnermann, A., Geuter, S., Sprenger, C., Finsterbusch, J. & Büchel, C. Interactions between brain and spinal cord mediate value effects in nocebo hyperalgesia. Science (80-.). 358, 105–108 (2017).

74. Chu, Y., Fricke, B. & Finsterbusch, J. NeuroImage Improving T2* - weighted human cortico-spinal acquisitions with a dedicated algorithm for region-wise shimming. Neuroimage 268, 119868 (2023).

75. Fricke, B. & Finsterbusch, J. A Shim Algorithm to Improve the Field Homogeneity and Image Quality in Cortico-Spinal fMRI. Proc. Int. Soc. Magn. Reson. Med. 28, 1216 (2020).

76. Jenkinson, M., Beckmann, C. F., Behrens, T. E. J., Woolrich, M. W. & Smith, S. M. FSL. Neuroimage 62, 782–790 (2012).

77. Jenkinson, M., Bannister, P., Brady, M. & Smith, S. Improved Optimization for the Robust and Accurate Linear Registration and Motion Correction of Brain Images. Neuroimage 17, 825–841 (2002).

78. Weber II, K. A., Chen, Y., Wang, X., Kahnt, T. & Parrish, T. B. Lateralization of cervical spinal cord activity during an isometric upper extremity motor task with functional magnetic resonance imaging. Neuroimage 125, 233–243 (2016).

79. De Leener, B. et al. PAM50: Unbiased multimodal template of the brainstem and spinal cord aligned with the ICBM152 space. Neuroimage 165, 170–179 (2018).

80. Glover, G. H., Li, T. Q. & Ress, D. Image-based method for retrospective correction of physiological motion effects in fMRI: RETROICOR. Magn. Reson. Med. 44, 162–167 (2000).

81. Chang, C., Cunningham, J. P. & Glover, G. H. Influence of heart rate on the BOLD signal: The cardiac response function. NeuroImage vol. 44 857–869 (2009).

82. Birn, R. M., Diamond, J. B., Smith, M. A. & Bandettini, P. A. Separating respiratory-variation-related fluctuations from neuronal-activity-related fluctuations in fMRI. Neuroimage 31, 1536–1548 (2006).

83. Grefkes, C., Eickhoff, S. B., Nowak, D. a, Dafotakis, M. & Fink, G. R. Dynamic intra-and interhemispheric interactions during unilateral and bilateral hand movements assessed with fMRI and DCM. Neuroimage 41, 1382–94 (2008).

84. Woolrich, M. W., Behrens, T. E. J., Beckmann, C. F., Jenkinson, M. & Smith, S. M. Multilevel linear modelling for FMRI group analysis using Bayesian inference. Neuroimage 21, 1732–1747 (2004).

85. Woolrich, M. W., Ripley, B. D., Brady, M. & Smith, S. M. Temporal autocorrelation in univariate linear modeling of FMRI data. Neuroimage 14, 1370–1386 (2001).

86. Beckmann, C. F., Jenkinson, M. & Smith, S. M. General multilevel linear modeling for group analysis in FMRI. Neuroimage 20, 1052–1063 (2003).

87. Woo, C.-W., Krishman, A. & Wager, T. D. Cluster-extent based thresholding in fMRI analyses: Pitfalls and recommendations. Neuroimage 91, 412–419 (2014).

88. Mayka, M. A., Corcos, D. M., Leurgans, S. E. & Vaillancourt, D. E. Three-dimensional locations and boundaries of motor and premotor cortices as defined by functional brain imaging: A meta-analysis. Neuroimage 31, 1453–1474 (2006).

89. RStudio Team. RStudio: Integrated Development Environment for R. (2021).

