## Supplementary material for "Ventral premotor cortex influences spinal cord activation during force generation"

| EV | X | Y | Z | Z-Value | Cluster size |
|---|---|---|---|---|---|
| **Mean** | 4 | -45 | -172 | 5.29 | 1690 |
| **Low** | 2 | -48 | -172 | 5.09 | 551 |
| **Medium** | 3 | -45 | -173 | 5.6 | 1027 |
| **High** | 2 | -47 | -170 | 5.36 | 2195 |

**Supp. Table S1 | FMRI derived task-related spinal activation.**
Group mean spinal BOLD activation. Listed are clusters with $P<0.001$, $Z>2.5$, cluster significance threshold of $P<0.05$. Peak coordinates and cluster sizes are given. EV explanatory variable.

| EV | Region | X | Y | Z | Z-Value | Cluster size |
|---|---|---|---|---|---|---|
| **Mean** | CL Postcentral G | -44 | -28 | 62 | 6.23 | 4685 |
| | CL Postc./Prec. G | -40 | -28 | 64 | 6.15 | |
| | CL Postc./Prec. G | -36 | -28 | 66 | 5.99 | |
| | | -18 | -10 | 16 | 5.91 | |
| | IL SMA | 4 | -8 | 56 | 6.39 | 2252 |
| | CL SMA | 0 | 0 | 46 | 5.96 | |
| | CL SMA | -4 | -6 | 52 | 5.95 | |
| | IL SMA | 10 | -8 | 62 | 5.93 | |
| | IL Supramarginal G | 52 | -40 | 48 | 6.05 | 2234 |
| | IL Parietal Operculum C | 60 | -32 | 26 | 5.81 | |
| | IL Superior Parietal L | 44 | -48 | 56 | 5.31 | |
| | IL Parietal Operculum C | 54 | -34 | 30 | 5.31 | |
| | IL Precentral G | 56 | 4 | 28 | 5.52 | 1271 |
| | IL Precentral G | 56 | 6 | 24 | 5.5 | |
| | IL Precentral G | 50 | 0 | 36 | 5.46 | |
| | | 36 | -6 | 34 | 5.37 | |
| | IL Frontal P | 36 | 36 | 30 | 5.53 | 390 |
| | IL Frontal P | 42 | 34 | 24 | 5.1 | |
| | | 28 | 20 | 24 | 4.76 | |
| | IL Middle Frontal G | 38 | 26 | 40 | 4.46 | |
| **Low** | CL Postcentral G | -46 | -28 | 58 | 5.95 | 2158 |
| | CL Prec. G/Postc. G. | -42 | -20 | 58 | 5.68 | |
| | CL Prec. G/Postc. G. | -34 | -30 | 70 | 5.57 | |
| | CL Prec. G/Postc. G. | -40 | -26 | 64 | 5.55 | |
| | IL SMA | 4 | -6 | 54 | 6.45 | 1526 |
| | CL SMA | -4 | -6 | 52 | 5.96 | |
| | IL SMA | 6 | -6 | 66 | 5.66 | |
| | CL SMA | -2 | -12 | 64 | 5.64 | |
| | IL Precentral G. | 54 | 4 | 28 | 5.99 | 1156 |
| | IL Precentral G. | 52 | -2 | 42 | 5.33 | |
| | IL Central Opercular C | 38 | 0 | 20 | 5.1 | |
| | | 18 | -4 | 16 | 4.98 | |
| | IL Lateral Occipital C | 36 | -68 | 56 | 5.16 | 791 |
| | IL Parietal Operculum C | 60 | -32 | 24 | 4.99 | |
| | IL Superior Parietal L | 42 | -54 | 60 | 4.88 | |
| | IL Supramarginal G | 56 | -40 | 48 | 4.86 | |
| | IL Middle Frontal G | 34 | 32 | 32 | 4.86 | 362 |
| | IL Middle Frontal G | 26 | 26 | 30 | 4.75 | |
| | IL Middle Frontal G | 38 | 24 | 36 | 4.66 | |
| | IL Middle Frontal G | 40 | 32 | 24 | 4.64 | |
| | CL Thalamus | -18 | -8 | 14 | 5.24 | 295 |
| | | -18 | -8 | 18 | 5.13 | |
| | | -22 | -6 | 16 | 5.12 | |
| | | -22 | -4 | 20 | 4.89 | |
| | CL Parietal Operculum C | -48 | -24 | 14 | 5.54 | 257 |
| | CL Supramarginal G | -58 | -24 | 26 | 4.2 | |
| | CL Central Opercular C | -62 | -18 | 14 | 4.01 | |
| | CL Central Opercular C | -44 | -20 | 24 | 3.62 | |
| **Medium** | CL Postcentral G | -46 | -26 | 60 | 6.07 | 2437 |
| | CL Prec. G/Postc. G. | -42 | -20 | 58 | 5.79 | |
| | CL Prec. G/Postc. G. | -36 | -28 | 66 | 5.78 | |
| | CL Prec. G/Postc. G. | -34 | -30 | 70 | 5.71 | |
| | IL SMA | 4 | -6 | 54 | 6.3 | 1427 |
| | CL SMA | -4 | -6 | 52 | 6.18 | |
| | CL SMA | -2 | -12 | 64 | 5.4 | |
| | IL SMA | 6 | -6 | 66 | 5.32 | |
| | IL Supramarginal G | 54 | -50 | 44 | 7.58 | 1032 |
| | IL Lateral Occipital C | 48 | -60 | 52 | 5.56 | |
| | IL Lateral Occipital C | 36 | -70 | 58 | 5.38 | |
| | IL Lateral Occipital C | 40 | -60 | 58 | 5.19 | |
| | IL Precentral G | 54 | 4 | 28 | 5.71 | 941 |
| | IL Precentral G | 52 | 0 | 40 | 5.11 | |
| | IL Precentral G | 46 | -6 | 46 | 4.85 | |
| | IL Middle Frontal G | 50 | 8 | 44 | 4.78 | |
| | IL Middle Frontal G | 42 | 34 | 30 | 6.41 | 292 |
| | IL Middle Frontal G | 34 | 36 | 30 | 4.9 | |
| | IL Middle Frontal G | 40 | 32 | 24 | 4.83 | |
| | IL Frontal P | 40 | 38 | 30 | 4.66 | |
| | CL Parietal Operculum C | -48 | -24 | 14 | 5.51 | 240 |
| | CL Central Opercular C | -60 | -20 | 16 | 4.17 | |
| | CL Central Opercular C | -44 | -20 | 24 | 3.67 | |
| | | -18 | -8 | 16 | 5.37 | 207 |
| | | -20 | -12 | 20 | 4.86 | |
| | | -16 | -16 | 12 | 4.78 | |
| | | -18 | -22 | 10 | 4.77 | |
| **High** | CL Postcentral G | -46 | -28 | 60 | 6.24 | 3848 |
| | CL Prec. G/Postc. G. | -40 | -28 | 64 | 6.03 | |
| | CL Prec. G/Postc. G. | -36 | -28 | 66 | 5.95 | |
| | CL Prec. G/Postc. G. | -42 | -20 | 58 | 5.92 | |
| | IL SMA | 4 | -8 | 56 | 6.23 | 2970 |
| | IL Middle Frontal G | 36 | 34 | 30 | 6.15 | |
| | CL SMA | -4 | -6 | 52 | 5.93 | |
| | IL SMA | 6 | -6 | 66 | 5.89 | |
| | IL Supramarginal G | 52 | -40 | 48 | 6.9 | 2234 |
| | IL Parietal Operculum C | 60 | -32 | 26 | 5.63 | |
| | IL Superior Parietal L | 44 | -48 | 56 | 5.31 | |
| | IL Supramarginal G | 52 | -34 | 36 | 5.27 | |
| | IL Precentral G | 54 | 4 | 28 | 5.66 | 1271 |
| | IL Precentral G | 50 | 0 | 36 | 5.56 | |
| | | 40 | 8 | 18 | 5.46 | |
| | IL Precentral G | 50 | 8 | 30 | 5.32 | |
| | CL Parietal Operculum C | -48 | -24 | 14 | 5.93 | 617 |
| | CL Postcentral G | -54 | -20 | 22 | 4.76 | |
| | CL Supramarginal G | -50 | -42 | 32 | 4.75 | |
| | CL Parietal Operculum C | -48 | -34 | 18 | 4.32 | |
| | | 20 | -4 | 16 | 6.57 | 209 |
| | | 18 | -2 | 22 | 4.85 | |
| | | 24 | 6 | 18 | 4.52 | |

**Supp. Table S2 | FMRI derived task-related cortical brain activation.**
Group mean cerebral BOLD activation. Up to 4 local maxima in a cluster are shown, CL=contralateral, IL=ipsilateral, G=Gyrus, C=Cortex, L=Lobule, SMA=Supplementary Motor Cortex, Prec. G=Precentral Gyrus, Postc. G=Postcentral Gyrus; Listed are all regions with *P*<0.001, Z>3.5, cluster significance threshold of *P*<0.05. EV explanatory variable. Identification of the regions with the Harvard-Oxford atlas.

| | M1 | | | SMA | | | PMV | | |
|---|---|---|---|---|---|---|---|---|---|
| **ID** | **X** | **Y** | **Z** | **X** | **Y** | **Z** | **X** | **Y** | **Z** |
| 1 | -40 | -28 | 50 | 0 | -10 | 56 | -56 | 5 | 16 |
| 2 | -43 | -26 | 48 | 0 | -12 | 67 | -52 | 2 | 38 |
| 3 | -29 | -25 | 65 | -2 | -4 | 50 | -59 | 6 | 21 |
| 4 | -35 | -30 | 48 | -9 | -12 | 64 | -54 | 1 | 34 |
| 5 | -29 | -30 | 54 | -2 | -15 | 60 | -52 | -9 | 42 |
| 6 | -29 | -36 | 54 | -2 | -8 | 65 | -56 | 8 | 28 |
| 7 | -33 | -34 | 50 | -4 | -14 | 68 | -58 | 6 | 34 |
| 8 | -40 | -19 | 64 | -4 | -6 | 49 | -52 | 0 | 26 |
| 9 | -36 | -34 | 58 | -4 | -21 | 59 | -56 | 0 | 32 |
| 10 | -30 | -34 | 55 | -2 | -8 | 64 | -56 | 13 | 27 |
| 11 | -28 | -32 | 50 | -8 | -9 | 54 | -54 | -3 | 39 |
| 12 | -28 | -22 | 67 | -1 | -6 | 54 | -50 | 8 | 38 |
| 13 | -36 | -34 | 54 | -7 | -21 | 50 | -60 | -8 | 22 |
| 14 | -36 | -23 | 58 | 0 | -14 | 58 | -56 | 0 | 22 |
| 15 | -36 | -29 | 52 | -4 | -22 | 64 | -52 | -4 | 22 |
| 16 | -32 | -28 | 52 | -8 | -24 | 54 | -53 | -5 | 42 |
| 17 | -26 | -35 | 63 | 0 | -12 | 54 | -58 | 5 | 30 |
| 18 | -38 | -24 | 44 | -13 | -10 | 69 | -60 | 0 | 22 |
| 19 | -38 | -28 | 59 | -2 | -10 | 64 | -44 | 3 | 39 |
| 20 | -34 | -26 | 59 | -4 | -12 | 70 | -53 | 2 | 30 |
| **Mean** | **-33.8** | **-28.9** | **55.2** | **-3.8** | **-12.5** | **59.7** | **-54.6** | **1.5** | **30.3** |

**Supp. Table S3 | Individual MNI peak coordinates of task-related brain activation**
Coordinates were derived from individual peak maxima for contralateral M1, SMA, PMV. Peak voxel activation was detected on the individual Z-maps and transformed to MNI. Peak coordinates were located using the mean EV.
